# Daytime broadcast spawning in cryptic *Pocillopora* species at Mo’orea, French Polynesia

**DOI:** 10.1101/2023.10.05.558016

**Authors:** P Harnay, AM Turner, SC Burgess, HM Putnam

## Abstract

Knowledge of coral gamete release is fundamental to reef ecology and evolution, but cryptic species complicate reproduction studies. In Mo’orea, French Polynesia, when corals were decimated by crown-of-thorns and a cyclone between 2007-2010, recruitment of six cryptic species of *Pocillopora spp.* primarily drove decadal reef recovery. Broadcast spawning for these genetically-verified species at Mo’orea is, however, undocumented in the scientific literature. We conducted *in situ* surveys over 58 days during September 2022-January 2023, October 2023-January 2024, and October 2024-December 2024. To identify the *Pocillopora spp.* we used molecular analysis of mtORF and PocHistone markers. *P. meandrina* and *P. grandis* spawned 2-3 days following the full moons of October-December. In contrast, *P. verrucosa* and *P. tuahiniensis* spawned at the new moons in November-December, with *P. verrucosa* spawning on lunar days 0-1, and *P. tuahiniensis* spawning on lunar days 1-3. Diel offset in the timing of spawning following sunrise was present in both moon phases indicating a temporal speciation barrier. *P. verrucosa* spawned ∼45 min after sunrise, whereas *P. tuahiniensis* spawned ∼73 min after sunrise. Similarly, *P. grandis* spawned ∼40 minutes after sunrise, whereas *P. meandrina* spawned ∼65 minutes after sunrise. Only one colony of *P. effusa* spawned (lunar day 3 new moon ∼06:30-06:50), requiring further confirmation of the timing for this low abundance species. Collectively, these first spawning reports provide an initial scientific documentation of *Pocillopora* spawning at the genetically-verified species level in Mo’orea, increasing capacity to study the essential process of coral reproduction for critical reef building species.

Data, Code, Image Archive: https://osf.io/5geqd/

**Résumé:** La connaissance de la libération des gamètes chez les coraux est fondamentale pour comprendre l’écologie et l’évolution des récifs, mais la présence d’espèces cryptiques complique les études sur la reproduction. À Mo’orea, en Polynésie française, lorsque les coraux ont été décimés par des étoiles de mer couronne d’épines et un cyclone entre 2007 et 2010, le recrutement de six espèces cryptiques de *Pocillopora spp*. a principalement contribué à la récupération des récifs au cours de la décennie suivante. Toutefois, la reproduction par émission synchronisée des gamètes (broadcast spawning) de ces espèces génétiquement vérifiées à Mo’orea n’a pas encore été documentée dans la littérature scientifique. Nous avons mené des observations in situ sur 58 jours entre septembre 2022 et janvier 2023, d’octobre 2023 à janvier 2024, puis d’octobre à décembre 2024. Pour identifier les espèces de *Pocillopora spp.*, nous avons utilisé une analyse moléculaire des marqueurs mtORF et PocHistone. *P. meandrina* et *P. grandis* ont pondu 2 à 3 jours après les pleines lunes d’octobre à décembre. En revanche, *P. verrucosa* et *P. tuahiniensis* ont pondu lors des nouvelles lunes en novembre et décembre, *P. verrucosa* aux jours lunaires 0 à 1, et *P. tuahiniensis* aux jours lunaires 1 à 3. Un décalage journalier dans le moment de la ponte après le lever du soleil a été observé pour les deux phases lunaires, indiquant une barrière temporelle de spéciation. *P. verrucosa* a pondu environ 45 minutes après le lever du soleil, tandis que *P. tuahiniensis* a pondu environ 73 minutes après. De manière similaire, *P. grandis* a pondu environ 40 minutes après le lever du soleil, alors que *P. meandrina* a pondu environ 65 minutes après. Une seule colonie de *P. effusa* a pondu (jour lunaire 3, nouvelle lune, entre ∼06:30 et 06:50), ce qui nécessite une confirmation supplémentaire du moment de reproduction pour cette espèce peu abondante. Collectivement, ces premières observations de ponte constituent une première documentation scientifique de la reproduction des *Pocillopora* au niveau des espèces génétiquement vérifiées à Mo’orea, augmentant la capacité à étudier ce processus essentiel de reproduction chez des espèces clés pour la construction des récifs.

## Introduction

Natural disturbances can trigger new ecological trajectories and states for a community (Knowlton 2015), with enduring ecological legacy effects (Cuddington 2011; Kopecky et al. 2023). However, natural disturbances such as storms and predator outbreaks (Kayal et al. 2012) have now combined with Anthropogenic disturbances, like increasing heat waves (Oliver et al. 2021), to degrade marine ecosystems, in particular coral reefs. Reef-building scleractinian corals are the ecosystem engineers that create 3D structure and provide vital resources to millions of people through critical ecosystem services, such as fisheries, food security, economic enhancement via tourism and biopharmaceuticals, indigenous cultural resources, and coastal protection from storms (Costanza et al. 2014). Consequently, natural maintenance of reef ecosystems relies on replenishment via coral reproduction and therefore the knowledge of coral reproductive timing and spawning is essential for basic coral biology and ecology, as well as coral selective breeding (van Oppen et al. 2015) and recruitment seeding (Doropoulos et al. 2019) in active reef restoration efforts.

Coral reef construction is based on the productive symbiotic relationship between the cnidarian host and symbiotic dinoflagellates (Symbiodiniaceae), which is susceptible to dysbiosis in response to relatively small environmental perturbations (e.g., 1°C over mean summer maxima (Lesser 2011)). Of particular concern in the Anthropocene is coral bleaching, or the coral-dinoflagellate symbiotic disruption driven by marine heatwaves that often results in substantial mortality of corals (Hughes, Kerry, and Simpson 2018) and cascading negative consequences for reef function (Courtney et al. 2018; Hughes et al. 2019). In French Polynesia, coral bleaching in 2002 resulted in >50% of corals surveyed showing bleaching in Mo’orea (Penin et al. 2007), and contributed to reduced coral cover (Pérez-Rosales et al. 2021). In Mo’orea in 2019, surveys of coral bleaching of the dominant genera revealed 89% of *Acropora* and 46% of *Pocillopora* had signs of bleaching and partial mortality (Speare et al. 2022). Additionally, another study of *Pocillopora* bleaching in April 2019 showed 72% of *Pocillopora* colonies bleached (Burgess et al. 2021). Further, bleaching occurred again in Mo’orea in March - May 2024 (MCR LTER pers comm). Therefore, without coral replenishment through reproduction, reef maintenance is threatened under current and future climate conditions in Mo’orea, and we also see evidence of these threats and degradation worldwide (Hughes et al. 2019; Reimer et al. 2024).

On multiple Polynesian islands, the genus *Pocillopora* has become the dominant space holder following a crown-of-thorns seastar (*Acanthaster planci*) outbreak in 2007 (Kayal et al. 2012) and a major cyclone, Oli, in 2010 (Pérez-Rosales et al. 2021). For example in Mo’orea, French Polynesia, when coral cover was reduced by nearly ∼90% due to crown-of-thorns outbreaks between 2007 and 2009 and cyclone Oli in 2010 (Holbrook et al. 2018), *Pocillopora spp.* were responsible for rapid reef recovery (Adjeroud, Penin, and Carroll 2007; Holbrook et al. 2018; Edmunds 2018). *Pocillopora spp.* are early colonizing species (Grigg and Maragos 1974) due to high fecundity (Sier and Olive 1994) and high recruitment (Adjeroud, Penin, and Carroll 2007). Further, with their complex branching morphologies, *Pocillopora spp.* provide extensive reef organism habitat (Stella, Jones, and Pratchett 2010; Pisapia et al. 2020). The Pocilloporid species diversity documented in Mo’orea and validated with genetics includes *P. meandrina*, *P. verrucosa, P. grandis*, *P. effusa, P. tuahiniensis (*Johnston and Burgess 2023; Johnston et al. 2022; Burgess et al. 2021; Burgess, Turner, and Johnston 2024), and *P. acuta* (primarily occurring on the fringing reefs; (Burgess, Turner, and Johnston 2024)). In light of this, it is essential that we quantify and describe the process of *Pocillopora* spp. spawning in French Polynesia, in order to understand and forecast the ecological dynamics of the reef in the future of increasing environmental disturbances.

A major impediment to quantifying and describing spawning patterns in corals, including *Pocillopora* spp., is the presence of cryptic species. Genetic identification to the species level in Mo’orea has clarified taxonomic differences in niche preference across reef zones and around the island (Burgess, Turner, and Johnston 2024), and also identified differential thermal sensitivity and mortality of *P. effusa* compared to other taxa in the 2019 coral bleaching event in Mo’orea (Burgess et al. 2021). Existing literature on reproduction in *Pocillopora* spp. (Baird, Guest, and Willis 2009; Schmidt-Roach et al. 2012) indicates they are hermaphroditic and primarily reproduce sexually via broadcast spawning (the release of both sperm and eggs into the water column). However, *P. damicornis*, and *P. acuta* can produce asexual larvae (Yeoh and Dai 2010; Combosch and Vollmer 2013) and reproduce sexually by brooding fertilized eggs internally, or by broadcast spawning (Ward 1992; Schmidt-Roach et al. 2014). For *Pocillopora spp.*, the release of sperm, eggs, and larvae (depending on the species) typically occurs after sunrise. While not all past studies conducted genetic species identification, there are reports (Table 1) of spawning near the full moon for morphologically identified *P. meandrina* in Australia (Schmidt-Roach et al. 2012) and Hawai i (Apprill et al. 2009; Schmidt-Roach et al. 2014), and for morphologically identified *P. grandis* (previously called *P. eydouxi*) in Japan (Hirose, Kinzie, and Hidaka 2001; Kinzie 1993; Hirose, Kinzie, and Hidaka 2000). In contrast, observations of morphologically identified *P. verrucosa* spawning have been made primarily during the new moon (but see (Schmidt-Roach et al. 2012)) in the Red Sea in Saudi Arabia, Sudan (Bouwmeester, Berumen, and Baird 2011; Bouwmeester et al. 2021), and Eilat (Shlesinger and Loya 1985), as well as in South Africa (Kruger and Schleyer 1998) and in Japan (Hirose, Kinzie, and Hidaka 2001; Kinzie 1993; Hirose, Kinzie, and Hidaka 2000; Baird et al. 2022). As lunar phase outliers, visually identified *P. meandrina* have been reported spawning on the new moons in Australia (Schmidt-Roach et al. 2012) and Taiwan (Lin and Nozawa 2017; Mulla et al. 2021), but these were not genetically confirmed.

**Table 1.** *Pocillopora* spp. spawning times reported from the literature based on observations or histological evidence.

| Reference | Location | Species | Identification | Evidence | Date | Moon | Lunar Day | Local Time |
| --- | --- | --- | --- | --- | --- | --- | --- | --- |
| (Glynn et al. 1991) | Caño Island, Costa Rica | *P. elegans* | Morphological | Histology | 1985-1989 | Full | NA | NA |
| (Glynn et al. 1991) | Pearl Islands, Panama | *P. elegans* | Morphological | Histology | 1984, 1985, 1989 | Full | NA | NA |
| (Glynn et al. 1991) | Uva Island, Panama | *P. elegans* | Morphological | Histology | 1984-1990 | Full | NA | NA |
| (Glynn et al. 1991) | Santa Cruz Island, Galapagos | *P. elegans* | Morphological | Histology | 1985-1989 | Full | NA | NA |
| (Kinzie 1993) | Okinawa, Japan | *P. grandis* (previously *P. eydouxi*) | Morphological | Spawning Observed | May - August 1992 | Full | Full Moon +1 to 5 | 08:00 – 10:50 |
| (Hirose, Kinzie, and Hidaka 2000) | Okinawa, Japan | *P. grandis* (previously *P. eydouxi*) | Morphological | Spawning Observed | June - July 1998 | Full | Few days after full moon | NA |
| (Schmidt-Roach et al. 2012) | Lizard Island, Australia | *P. grandis* (previously *P. eydouxi*) | Morphological | Spawning Observed | 11-12 November 2011 | Full | Full moon +1-2 | NA |
| (Fiene-Severns 1998) | Molokini, Hawaiï | *P. meandrina* | Morphological | Spawning Observed | April-May; 1991, 1994-1998 | Full | Full moon +2-5 | 07:20 |
| (Schmidt-Roach et al. 2012) | Lizard Island, Australia | *P. meandrina* | Morphological | Spawning Observed | 11-12 November 2011 | Full | Full moon +1-2 | 06:25 |
| (Fadlallah 1985) | Yanbu, Saudi Arabia | *P. verrucosa* | Morphological | Histology | May June | New | NA | NA |
| (Shlesinger and Loya 1985) | Eilat, Israel | *P. verrucosa* | Morphological | Spawning Observed | July - August 1981-1982 | New | NA | NA |
| (Kinzie 1993) | Okinawa, Japan | *P. verrucosa* | Morphological | Spawning Observed | May – July 1992 | New | New Moon +1 to 6 | 07:00 – 07:11 |
| (Sier and Olive 1994) | Maldives | *P. verrucosa* | Morphological | Histology | March - April 1991 | NA | NA | NA |
| (Tomascik 1997) | Karimunjawa Islands, Indonesia | *P. verrucosa* | Morphological | not provided | October - November 1995 | NA | NA | NA |
| (Kruger and Schleyer 1998) | South Africa | *P. verrucosa* | Morphological | Histology | January-February 1993 | New | NA | NA |
| (Hirose, Kinzie, and Hidaka 2000) | Okinawa, Japan | *P. verrucosa* | Morphological | Spawning Observed | June - July 1998 | New | Few days after new moon | NA |
| (Romatzki, 2008) | Sulawesi, Indonesia | *P. verrucosa* | Morphological | Histology | March, August 2006-2007 | New | NA | NA |
| (Bouwmeester, Berumen, and Baird 2011) | Thuwal, Saudi Arabia | *P. verrucosa* (now *P. favosa*) | Morphological | Spawning Observed | 31 May 2011 | New | New moon - 1 | 8:40 - 9:15 |
| (Schmidt-Roach et al. 2012) | Lizard Island, Australia | *P. verrucosa* | Morphological | Spawning Observed | 11-12 November 2011 | Full | Full moon +1-2 | NA |
| (Suzuki 2012) | Ryukyu Islands, Japan | *P. verrucosa* | Morphological | Spawning Observed | 18 June 2011 | Full | NA | 09:00 |
| (Bouwmeester et al. 2015) | Dreams Beach Reef, Saudi Arabia | *P. verrucosa* (now *P. favosa*) | Morphological | Spawning Observed | May 2011, May 2012 | New | New moon - 1 | 8:40 - 9:20 |
| (Lin and Nozawa 2017) | Lyudao, Taiwan | *P. verrucosa* | Morphological | Spawning Observed | April - June 2011 - 2015 | Full | Full +1 +2 +3 | 09:50–11:00 |
| (Mulla et al. 2021) | Lyudao, Taiwan | *P. verrucosa* | Morphological | Spawning Observed | 21 April 2019 | Full | Full +2 +3 | 9:30 and 10:00 |
| This study | Mo'orea, French Polynesia | *P. meandrina* | Genetic | Spawning Observed | October 2023, November 2022, December 2022 | Full | Full +2 +3 | 6:16-6:38 |
| This study | Mo'orea,<br>French<br>Polynesia | *P. grandis* | Genetic | Spawning<br>Observed | November,<br>December<br>2024 | Full | Full +2<br>+3 | 5:55-6:12 |
| This study | Mo'orea,<br>French<br>Polynesia | *P. verrucosa* | Genetic | Spawning<br>Observed | November,<br>December<br>2023 | New | New<br>+1 +2 | 5:56-6:25 |
| This study | Mo'orea,<br>French<br>Polynesia | *P. tuahiniensis* | Genetic | Spawning<br>Observed | November,<br>December<br>2023 | New | New<br>+1 +2<br>+3 | 6:19-6:58 |
| This study | Mo'orea,<br>French<br>Polynesia | *P. effusa* | Genetic | Spawning<br>Observed | November<br>2023 | New | New<br>+3 | 06:30-06:50 |

**Table 2.**
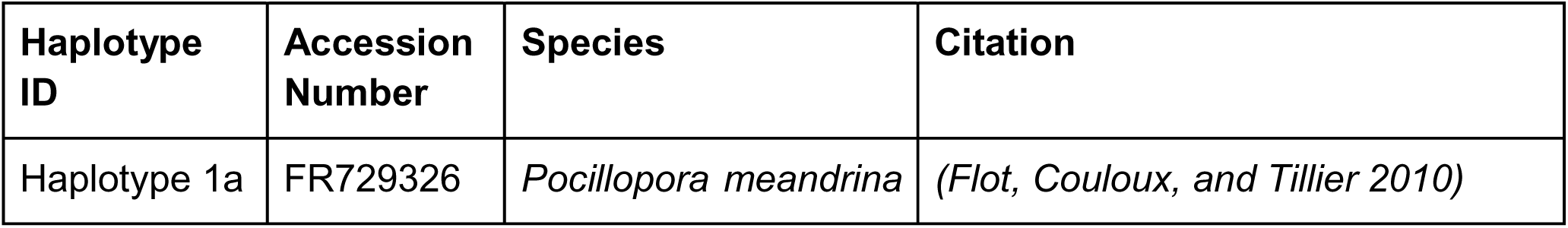
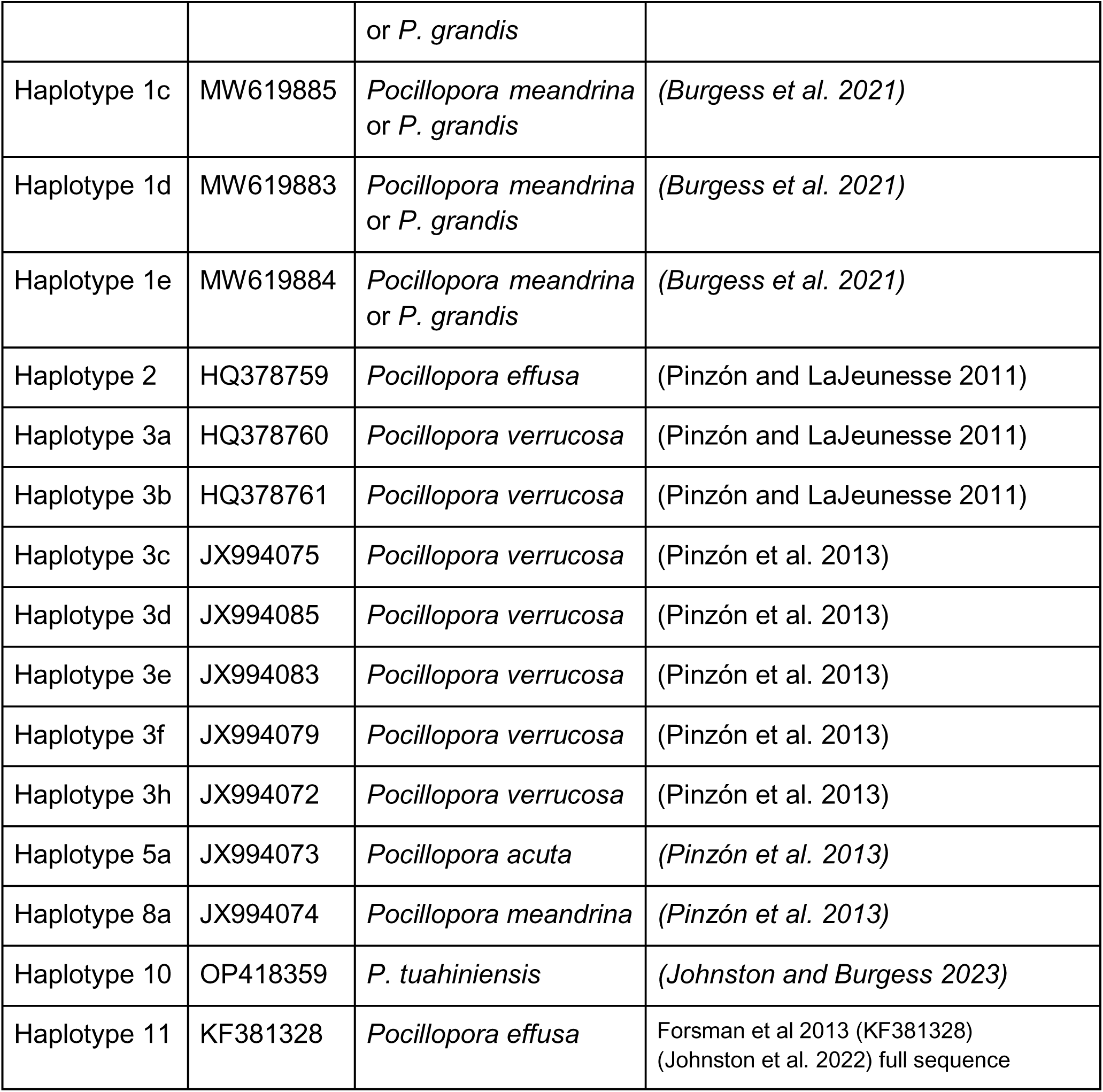
Reference mtORF sequences and their accession numbers used for sequence alignment and tree building. Raw sequence data, fasta files, and alignment are archived at https://osf.io/5geqd/ (Harnay et al. 2023)

| Haplotype ID | Accession Number | Species | Citation |
| --- | --- | --- | --- |
| Haplotype 1a | FR729326 | *Pocillopora meandrina* or *P. grandis* | (Flot, Couloux, and Tillier 2010) |
| Haplotype 1c | MW619885 | *Pocillopora meandrina*<br>or *P. grandis* | (Burgess et al. 2021) |
| Haplotype 1d | MW619883 | *Pocillopora meandrina*<br>or *P. grandis* | (Burgess et al. 2021) |
| Haplotype 1e | MW619884 | *Pocillopora meandrina*<br>or *P. grandis* | (Burgess et al. 2021) |
| Haplotype 2 | HQ378759 | *Pocillopora effusa* | (Pinzón and LaJeunesse 2011) |
| Haplotype 3a | HQ378760 | *Pocillopora verrucosa* | (Pinzón and LaJeunesse 2011) |
| Haplotype 3b | HQ378761 | *Pocillopora verrucosa* | (Pinzón and LaJeunesse 2011) |
| Haplotype 3c | JX994075 | *Pocillopora verrucosa* | (Pinzón et al. 2013) |
| Haplotype 3d | JX994085 | *Pocillopora verrucosa* | (Pinzón et al. 2013) |
| Haplotype 3e | JX994083 | *Pocillopora verrucosa* | (Pinzón et al. 2013) |
| Haplotype 3f | JX994079 | *Pocillopora verrucosa* | (Pinzón et al. 2013) |
| Haplotype 3h | JX994072 | *Pocillopora verrucosa* | (Pinzón et al. 2013) |
| Haplotype 5a | JX994073 | *Pocillopora acuta* | (Pinzón et al. 2013) |
| Haplotype 8a | JX994074 | *Pocillopora meandrina* | (Pinzón et al. 2013) |
| Haplotype 10 | OP418359 | *P. tuahiniensis* | (Johnston and Burgess 2023) |
| Haplotype 11 | KF381328 | *Pocillopora effusa* | Forsman et al 2013 (KF381328)<br>(Johnston et al. 2022) full sequence |

Multiple recruitment studies in Mo’orea indicate the density of Pocilloporidae recruits peaks between December - April each year (Gleason 1996; Adjeroud, Penin, and Carroll 2007; Edmunds, Leichter, and Adjeroud 2010). Therefore, we focused our examination of *Pocillopora spp.* spawning in the months of October - January. In this study, we conducted *in situ* surveys of *Pocillopora* spp. spawning over a total of 58 days in the lagoon of the north shore of Mo’orea French Polynesia (Table S1). On each day, observations lasted for ∼2-3 hours following sunrise for several days around the full and new moons (where logistics and conditions allowed) for the months of September - January in 2022-2023, October - January in 2023-2024, and October - December in 2024. Corals observed spawning were genetically identified to species (Table S2). Collectively, this allowed for clear documentation of the lunar and diel timing of spawning of *P. grandis, P. meandrina*, *P. verrucosa*, and *P. tuahiniensis*, as well as a possible indication of *P. effusa* timing.

## Methods

In order to describe the lunar and diel spawning times to the species level, we completed *in situ* visual and photographic/video documentation of *Pocillopora* spp. spawning and confirmed successful embryo development, as well as analyzed samples for species level genetic identification.

### Spawning observations and sample collection

For this work, we followed the lunar calendar dates set by the United States Astronomical Applications Department (https://aa.usno.navy.mil/data/MoonPhases). The spawning observation site was located on the north shore lagoon reef in Mo’orea (- 17.475757, -149.808062) covering approximately an area of 400m^2^, which was selected for high Pocilloporidae cover (Edmunds 2022) and capacity for safe low light small boat transit from the UC Richard B. Gump South Pacific Research Station. All observations were made while snorkeling in ∼3m, with a minimum of two observers in the water. Observers were positioned to simultaneously visualize multiple morphologies with the goal of observing five common lagoon species (*P. effusa*, *P. grandis, P. meandrina*, *P. verrucosa*, and *P. tuahiniensis*) (Burgess, Turner, and Johnston 2024). Coral spawning observations were documented as follows: (1) photographs of each colony were taken during or following spawning (Canon G7X MarkII); (2) colonies were tagged and geolocated via GPS (Garmin GPSMAP 64); (3) biopsies were collected from each tagged colony in sterile Whirlpaks (Fig. 1-4); and (4) where possible, gametes were collected from the water next to the colonies using plastic bottles. After collecting eggs and sperm from different colonies, the bottles used for collection were poured into larger reservoirs (∼5L bottles) to facilitate genetic mixing, as well as fertilization as soon as possible directly on a boat. Samples were returned to the Gump South Pacific Research Station and biopsies from the tagged colonies were preserved in DNA/RNA Shield stabilization solution (Zymo Research Corporation, Cat # R1100). To confirm successful fertilization occurred, collected gametes were viewed with a compound microscope (LEICA DM500)(e.g., Fig. 1).

**Figure 1.**
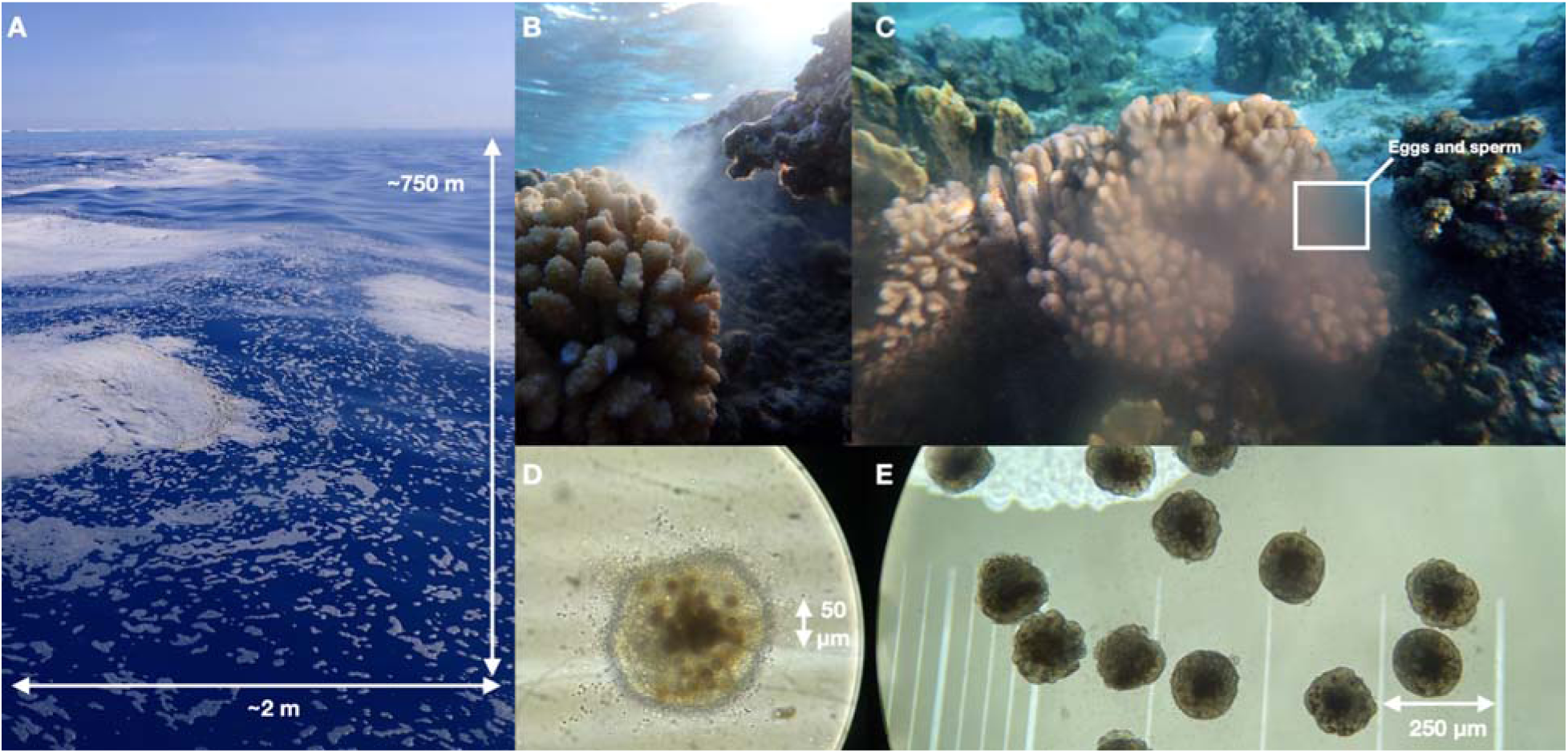
**A)** Slick of sperm and egg on the surface of the water in the lagoon was observed after *Pocillopora meandrina* spawning, with the image showing ∼ 750m distance between the fore reef and the observer. **B)** and **C)** Spawning of *P. meandrina* colonies in the lagoon. **D)** Brightfield microscopy image of eggs (brown circle) and sperm (gray dots) surrounding the egg. **E)** Embryos at a multi-cell stage were visible at ∼4 hours post spawning. More images and videos are archived here: https://osf.io/5geqd/ (Harnay et al. 2023)

### *Pocillopora* species genetic identification

The use of mtORF sequencing and PocHistone PCR plus *Xho*I digestion for Restriction Fragment Length Polymorphism (RFLP) has previously been validated as suitable species-level markers based on multiple genome-wide datasets and analyses (Johnston, Forsman, and Toonen 2018; Johnston, Cunning, and Burgess 2022). Through these methods, the following *Pocillopora* species have been defined as: *P. meandrina* = mtORF haplotype 1 followed by a single *Xho*I digestion RFLP band of the PocHistone amplification product, or mtORF haplotype 8a and haplotype 9; *P. grandis* = mtORF haplotype 1 followed two or three *Xho*I digestion RFLP bands of the PocHistone amplification product; *P. tuahiniensis* = mtORF haplotype 10; *P. verrucosa* = mtORF haplotype 3; and *P. effusa* = mtORF haplotypes 2 and 11.

In order to determine the species of *Pocillopora* spawning, DNA was extracted from each sample using the Quick-DNA Miniprep Plus Kit (Zymo Research Corporation Cat # D4069) or Chelex 100 Resin (Bio-Rad Cat #142-1253). For the Quick-DNA extractions, DNA was extracted according to the manufacturer’s instructions for samples stored in DNA/RNA Shield, including the addition of Proteinase K (20mg mL^-1^). For the Chelex extractions, 10 μL of DNA Shield was placed into 100 μL of 10% Chelex solution and incubated at 55°C for 60 minutes followed by 95°C for 20 minutes. The sample was centrifuged at 4,000 rpm for 2 minutes and the supernatant DNA moved to a new tube.

To identify *Pocillopora* spp. we amplified the mitochondrial open reading frame (mtORF) region using the primers of Flot et al. (Flot et al. 2008) FatP6.1 (5′-TTTGGGSATTCGTTTAGCAG-3′) and RORF (5′-SCCAATATGTTAAACASCATGTCA-3′). For the Zymo Quick-DNA extractions, PCR mixes contained 12.5 µL of EmeraldAmp GT PCR Master Mix (TaKaRa Bio USA Inc. Cat # RR310B), 0.3 µL of each of the forward and reverse primers (10 µM), 1 µL of template DNA, and 10.9 µL of deionized water to 25 µL final volume per sample. For the Chelex extractions, PCR mixes contained 5 µL BioMix Red (Bioline Ltd., London, UK Cat # BIO-25006), 0.13 µL of each of the forward and reverse primers (10 µM), 0.1 µL Recombinant Albumin (New England Biolabs Cat # B9200S), 1 µL of template DNA, and 3.64 µL of deionized water to 10 µL final volume per sample. Sample DNAs were amplified with the polymerase chain reaction (PCR) profile of a single denaturation step of 60s at 94 °C, followed by 30s at 94 °C for denaturation, 30s at 53 °C for annealing, and extension of 75s at 72 °C for 30 cycles (2022 samples) or 40 cycles (2023 samples); thermocycling was followed by final incubation at 72 °C for 5 min. The products were then assessed with a 1.5% or 2% agarose gel in TAE for 45 mins at 100 V or 30 mins at 90 V to verify bands of ∼1000bp.

The PCR products from the Quick-DNA extractions (2022 samples) were cleaned with Qiagen MinElute PCR Purification Kit (Qiagen Cat # 28004) according to the manufacturer’s instructions. The PCR products from the Chelex extractions (2023 samples) were cleaned with Exo-CIP Rapid PCR Cleanup (New England Biolabs Cat # E1050L) according to the manufacturer’s instructions. Sanger sequencing was completed with forward primers at the URI Genomics and Sequencing center using Applied Biosystems BigDye Terminator v3.1 chemistry (2022 samples) and at the Florida State University Core Facility using the Applied Biosystems 3730 Genetic Analyzer with Capillary Electrophoresis (2023 samples). Sequences were aligned using Geneious Alignment in GENEIOUS PRIME 2020.2.4. Pairwise alignment of 723bp from the 72 sample sequences along with the 16 reference sequences (https://osf.io/5geqd/) was completed using Clustal Omega (Sievers et al. 2011) in GENEIOUS PRIME 2020.2.4. Tree building was conducted with a Jukes-Cantor genetic distance model (Jukes and Cantor 1969) and no outgroup. The species tree was generated via Neighbor-Joining estimation (Saitou and Nei 1987) with the Neighbor-Joining function from the package phangorn (Schliep 2011) in R (Team 2013), followed by Ward’s hierarchical agglomerative clustering method (ward.D; (Murtagh and Legendre 2014). Raw sequence data and alignment fasta files are archived here: https://osf.io/5geqd/ (Harnay et al. 2023).

When mtORF sequencing identified samples as Haplotype 1 (containing both *P. grandis* and *P. meandrina*), DNAs were also PCR amplified with the histone 3 region using the primers of Johnston et al. (Johnston, Forsman, and Toonen 2018) (PocHistoneF: 5′-ATTCAGTCTCACTCACTCACTCAC-3′ and PocHistoneR: 5′-TATCTTCGAACAGACCCACCAAAT-3′), including a previously identified *P. grandis* as a positive control. PCR mixes were made following the methods above (but with PocHistone primers). Thermal cycling included a denaturation step of 60 s at 94 °C, followed by 30 cycles (2022 samples) or 40 cycles (2023 samples) of 30 s at 94 °C, 30 s at 53 °C, and 60 s at 72 °C; and an elongation step at 72 °C for 5 min. PCR products were then used for Restriction Fragment Length Polymorphism (RFLP) with *Xho*I (New England BioLabs, NEB R0146S) in rCutSmart buffer (New England BioLabs, NEB B6004S) according to Johnston et al. (Johnston, Forsman, and Toonen 2018) (with a volume exception as follows for 2023 samples; 5 µL of PCR product were digested with 0.5µL of *Xho*I restriction enzyme, 0.8 µL of 10X CutSmart buffer, and 2.7µL of deionized water, for a final volume of 9µL). Restriction digest products were viewed on a 2% agarose gel in 1X TAE after running for ∼90-110 min at 70V with a GeneRuler 100 bp DNA Ladder (Thermo Fisher Cat SM0241) or Quick-Load Purple 100 bp DNA Ladder (New England Biolabs NEB N0467S). A single band at 669 bp indicates the species is *P. meandrina*, while two bands at 382 and 287 bp indicate the species identification is *P. grandis* (Johnston, Forsman, and Toonen 2018). Colonies observed spawning in 2024 were previously tagged and extracted with the Chelex methods as described above, amplified with the 2022 methods and sequenced as follows. The PCR product was cleaned by mixing 3µl of 3M sodium acetate and 3x volume of ice cold 100% ethanol with the PCR product and holding it in the -80°C freezer for 30 min. The samples were then spun for 30 minutes at 12,000 rpm, the supernatant discarded, and the DNA pellet rinsed with ice cold 70% ethanol, spun for 5 minutes at 12,000 rpm, and left to dry at room temperature until all ethanol evaporated. Sanger sequencing was completed with forward primers at the URI Genomics and Sequencing center using Applied Biosystems BigDye Terminator v3.1 chemistry. Raw sequence data, alignment fasta files, and gel images are archived here: https://osf.io/5geqd/ (Harnay et al. 2023)

## Results

### Full Moon Observations

In our observations of full moon spawning, *Pocillopora meandrina* and *P. grandis* were observed releasing eggs and sperm (Table S1-2; Fig. 2). We witnessed gamete release in *P. grandis* on lunar days 2 and 3 after the full moon in the months of November and December in 2024 (Fig. 4). Spawning occurred between 36 and 42 minutes after sunrise (Fig. 5). Observations of *P. meandrina* conducted between 2022 and 2024 revealed spawning on lunar day 1 to 3 after the full moon between October and December (Fig. 4), with the largest events occurring in December. Gamete release occurred later in the morning after *P. grandis*, between 59 and 77 minutes after sunrise (Fig. 5).

**Figure 2.**
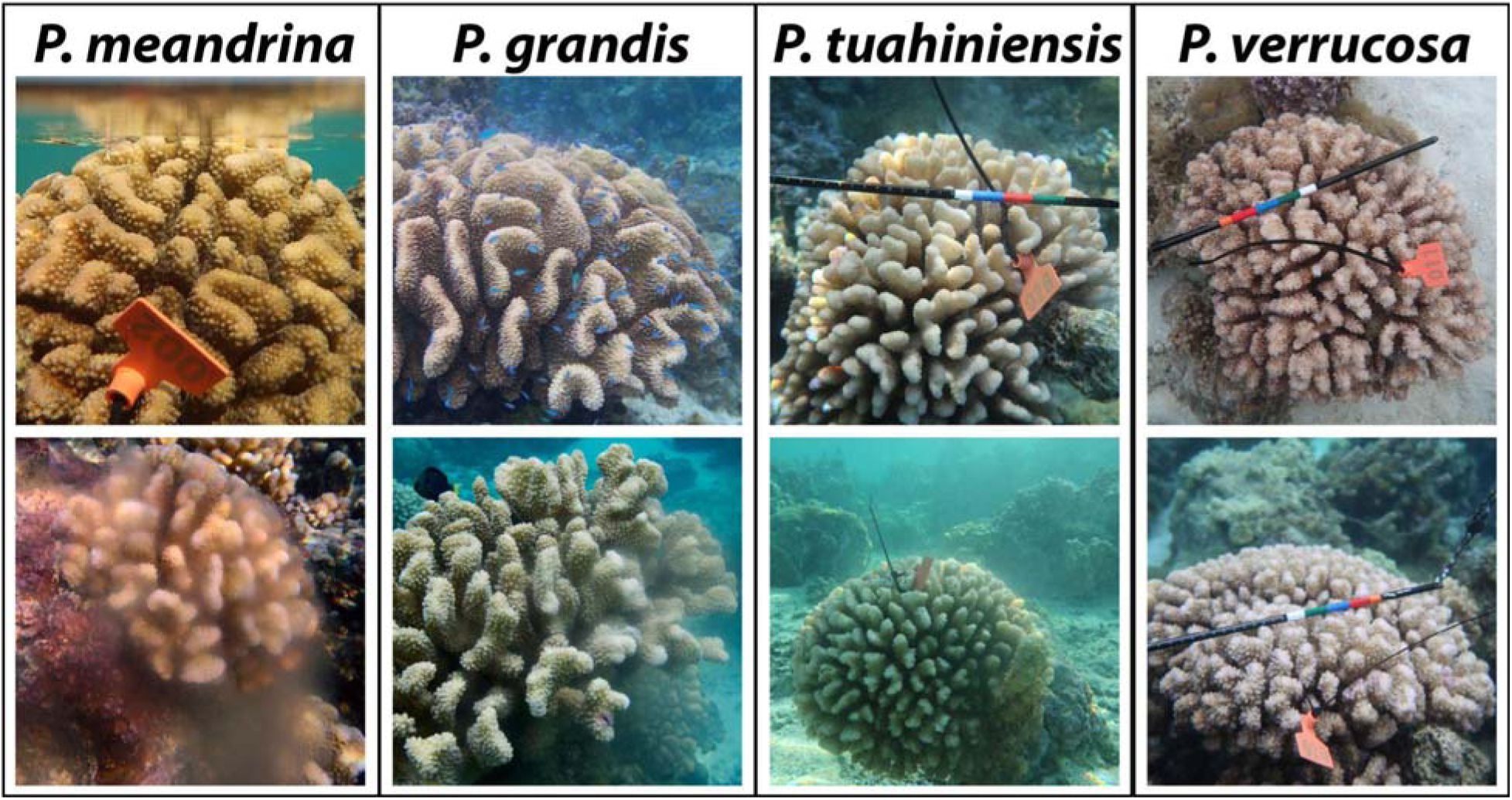
Examples of *Pocillopora* colonies that were tagged and observed spawning eggs and sperm in the lagoon.

**Figure 3.**
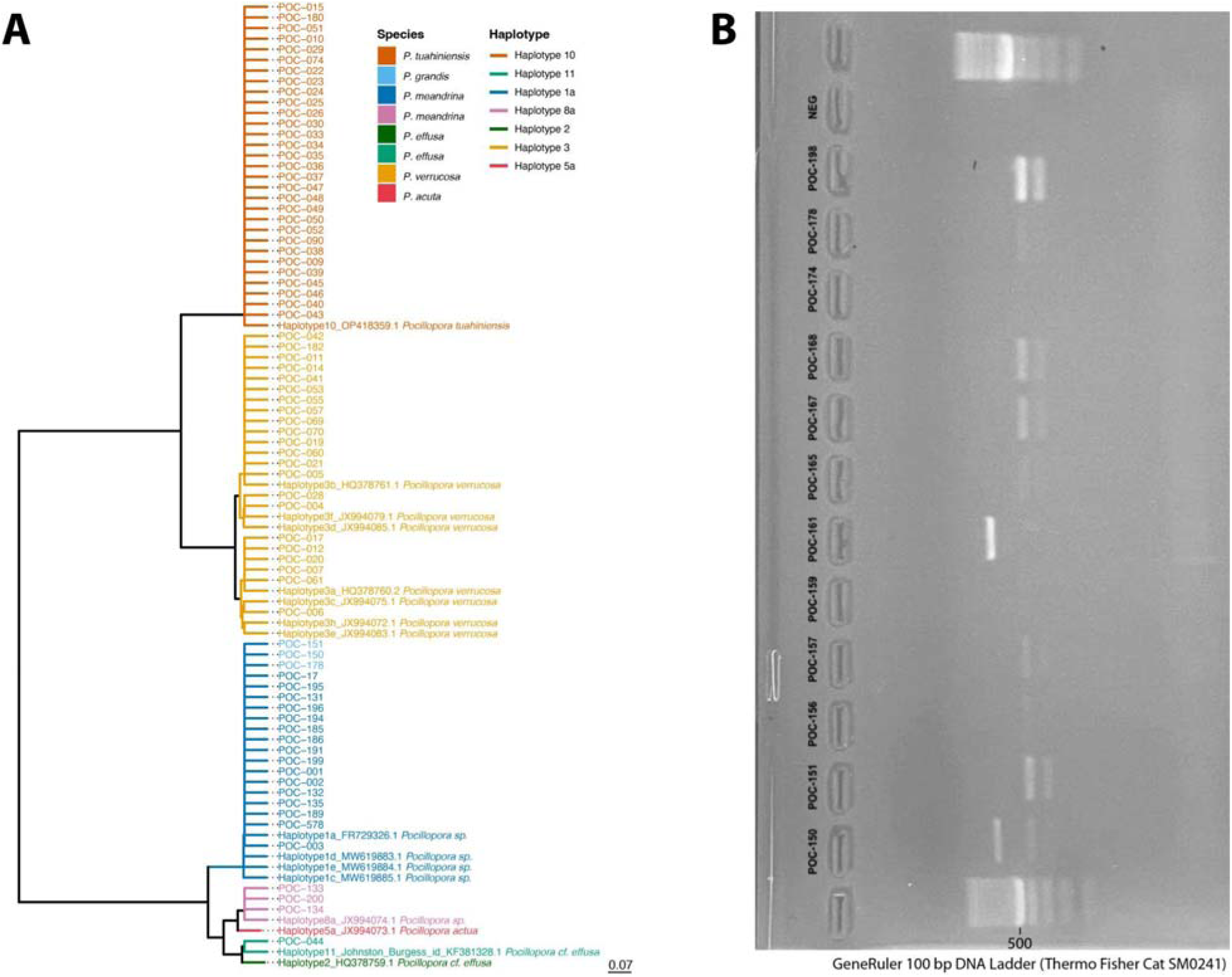
Haplotype identification via mtORF for spawning colonies in 2022, 2023, and 2024 generated from hierarchical clustering of Jukes-Cantor genetic distance (Jukes and Cantor 1969) from 75 collected samples, 16 reference samples, and no outgroup. The gene tree is based on mtORF to visualize species delimitation and does not reflect exact phylogeny (see (Johnston, Cunning, and Burgess 2022)). Species colors are added based on unambiguous mtORF haplotypes, or following unambiguous PocHistone RFLP results. Raw sequence data, alignments, gel images, and reference fasta files are archived here: https://osf.io/5geqd/ (Harnay et al. 2023).

**Figure 4.**
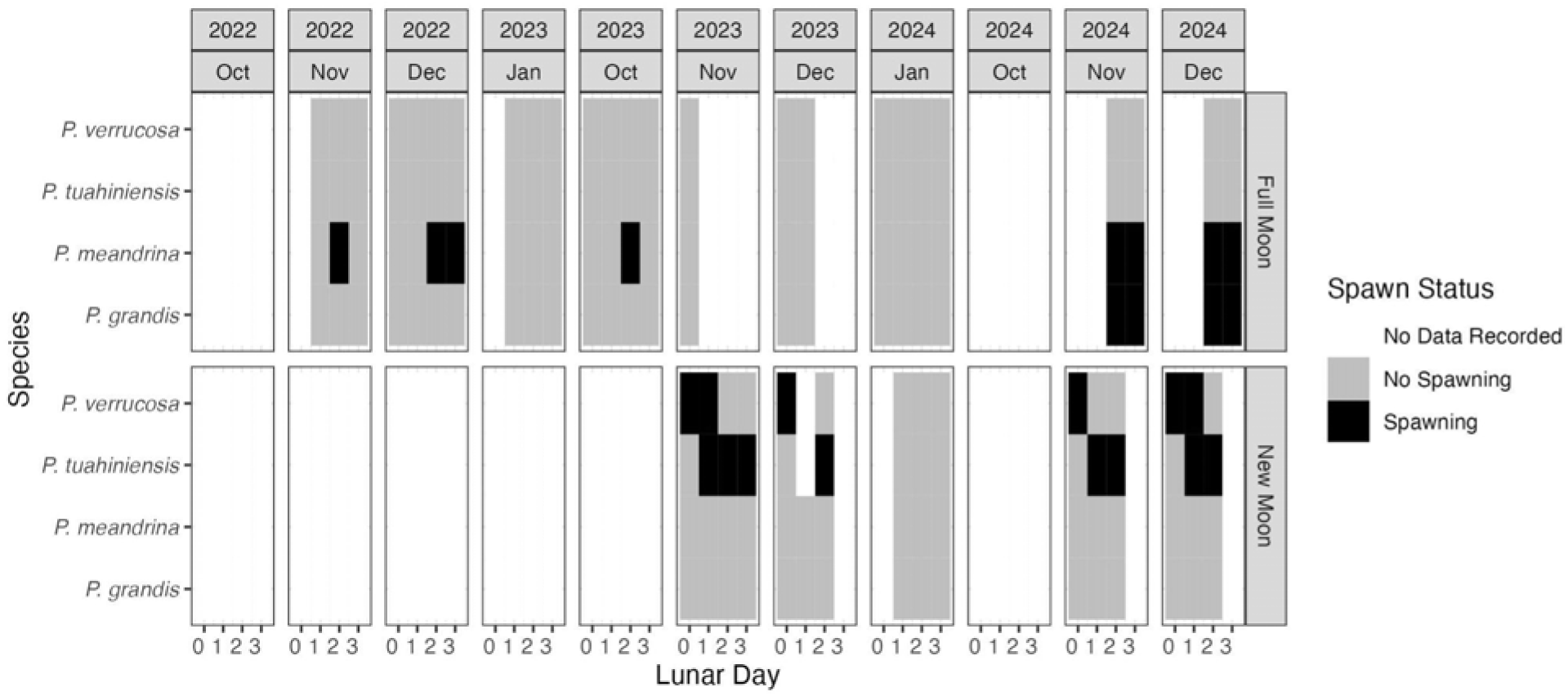
Lunar timing of spawning on day 0 (full or new moon) and the following 3 days during the months surveyed from October 2022 to December 2024.

**Figure 5.**
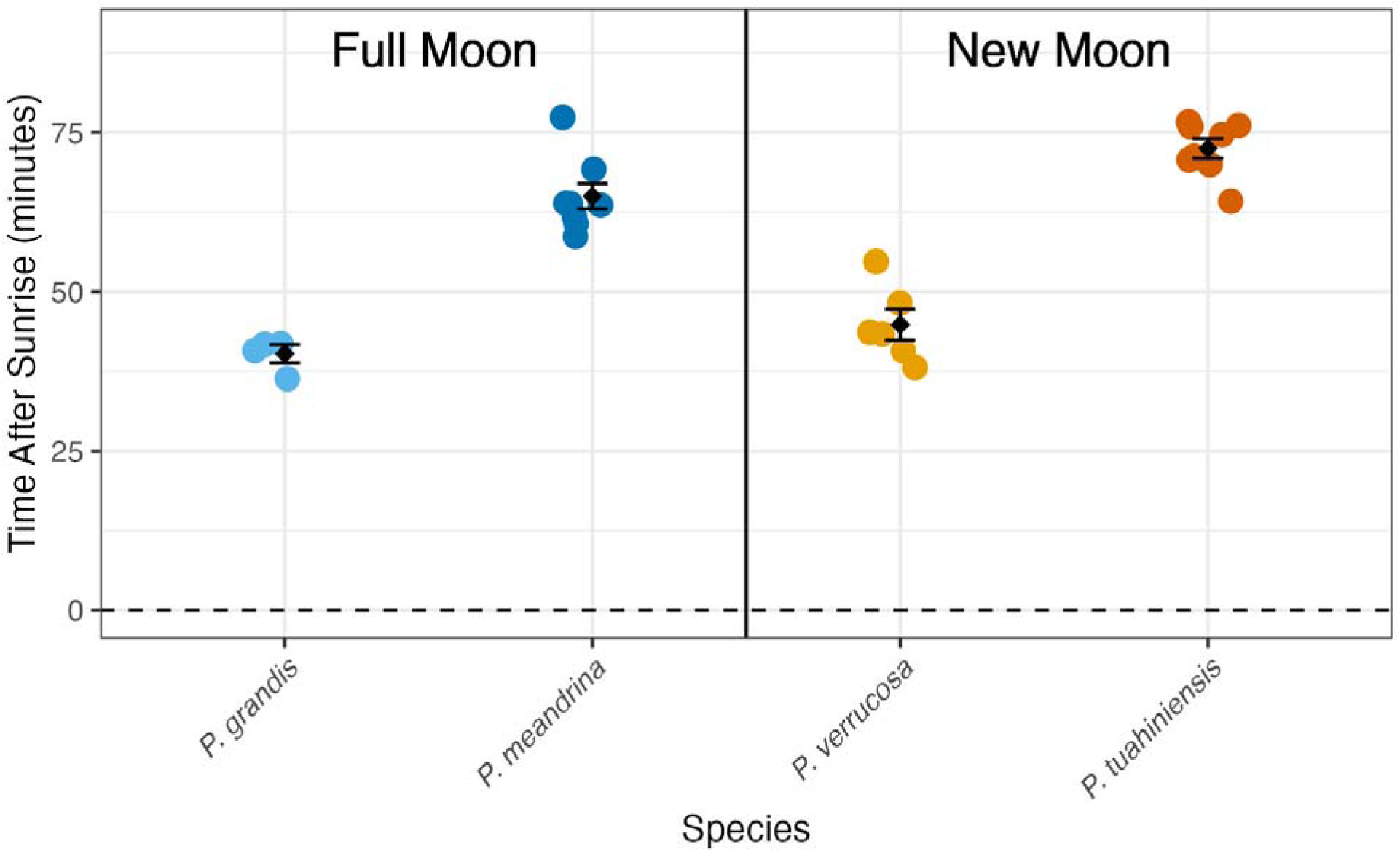
Diel timing of spawning (minutes after sunrise) for *P. grandis* and *P. meandrina* around the full moon and for *P. verrucosa* and *P. tuahiniensis* around the new moon. Colored circles indicate an individual coral, while black diamonds indicate the mean value ± standard error of the mean.

### New Moon Observations

In our new moon spawning observations, colonies were predominantly identified as *P. verrucosa* and *P. tuahiniensis* (Fig. 3). *P. verrucosa* spawning was observed spawning in November and December 2023 and 2024, on lunar days 0 and +1 (Fig. 4), between 38 and 55 minutes after sunrise (Fig. 5). *Pocillopora tuahiniensis*, spawning was observed in 64 to 77 minutes after sunrise (Fig. 5), also in November and December of 2023 and 2024, but from day +1 to +3 (Fig. 4).

The least abundant species detected by mtORF sequencing during the new moon (Table S2) was a single colony of *P. effusa* identified as Haplotype 11 (Johnston et al. 2022). The *P. effusa* observation occurred in November 2023 sampling window of ∼06:30 - 06:50 on the new moon lunar day 3. This species was not included in the figures because of the small sample size (n=1), and therefore we do not have enough observations for this species to demonstrate a clear spawning pattern. See repository for image; https://osf.io/5geqd/).

### Larvae contain zooxanthellae and development

For all *Pocillopora* species we observed, once spawning commenced the water quickly became cloudy with eggs and sperm (e.g., Fig. 1A, B, C). Eggs appeared to be neutrally or negatively buoyant in the collection bottles and in the water the gametes were rapidly dispersed by the waves and current. Eggs (∼90µm) and developing embryos (∼120µm) that were observed at ∼10:00 via compound microscopy contained zooxanthellae. By 14:00 (∼7.5 hours post fertilization, hpf) larvae were actively swimming.

## Discussion

Multiple recruitment studies in Mo’orea indicate the density of Pocilloporidae recruits peaks between December - April each year (Gleason 1996; Adjeroud, Penin, and Carroll 2007; Edmunds, Leichter, and Adjeroud 2010). Therefore, we focused our examination of *Pocillopora* spp. spawning between the month of October and January during the years 2022, 2023 and 2024, resulting in the clear description of the timing of spawning for *P. meandrina*, *P. verrucosa, P. tuahiniensis*, *P. grandis* and the start of documentation for *P. effusa*.

### Pocillopora meandrina

During our observations of *P. meandrina*, gametes were released early in the morning at ∼06:15-06:40, which was ∼ 60-85 minutes after sunrise local time. This information aligned with the range of (morphologically identified) *Pocillopora* spp. spawning in the literature of ∼45-180 minutes after sunrise between different locations around the world, such as Australia (∼45 minutes after sunrise (Schmidt-Roach et al. 2012)), Okinawa, Japan (∼95 to 120 minutes after sunrise (Hirose, Kinzie, and Hidaka 2001; Kinzie 1993; Hirose, Kinzie, and Hidaka 2000; Baird et al. 2022)), Hawai i (∼120 minutes after sunrise (Schmidt-Roach et al. 2014)), and the Red Sea (∼180 minutes after sunrise (Bouwmeester, Berumen, and Baird 2011; Bouwmeester et al. 2021)). We documented split-spawning (spawning in multiple consecutive months) in *P. meandrina*, with one colony spawning November 10th (06:32, full moon lunar day 3) and reef-wide spawning December 9th (06:16-06:32; full moon lunar day 2) and December 10th (06:21-06:35; full moon lunar day 3), which could be seen as surface slicks (Fig. 1A). Split spawning across multiple months is in line with spawning observations of (morphologically identified) *P. meandrina* in Hawai i (Fiene-Severns 1998; Kolinski and Cox 2003), and the congeners (morphologically identified) of *P. verrucosa* and *P. grandis* (previously *P. eydouxi*) in Japan (Kinzie 1993).

In December 2022, most of the *P. meandrina* colonies observed had synchronized spawning, based on the observation that on December 9 (Full+2), from 6:16 to 6:32, a large number of *P. meandrina* colonies were observed spawning both eggs and sperm turning the water of the lagoon cloudy and producing surface slicks (Fig. 1A). Personal communications from other scientists in the region confirmed that *Pocillopora* spp. were spawning on forereef sites in Tetiaroa Dec 9th ∼06:30 (Camille Gâche, Pers Comm) and Rangiroa Dec 9th ∼06:20 (Camille Leonard, Pers Comm). On December 10th 2022 (Full+3), from 6:21 to 6:35, we observed a second day of synchronized spawning, but with a lower abundance of colonies releasing. Our results corroborate those from Japan (Kinzie 1993) and Hawai i (Fiene-Severns 1998; Kolinski and Cox 2003) that *Pocillopora* spp. can spawn across multiple days within a month.

Across our observation period of October 2023 - January 2024, we were unable to access the observation site for all months and all desired days around the full moon (−2 to +5) when we hypothesized *P. meandrina* spawning could be occurring based on our 2022 observations, the Pocilloporid recruitment window of November to March from Mo’orea (Adjeroud, Penin, and Carroll 2007; Edmunds, Leichter, and Adjeroud 2010), and from the hypothesized timing of *Pocillopora* spp. spawning from emerging histological data (Putnam Unpublished Data). Given that pocilloporid recruitment continues through March (Adjeroud, Penin, and Carroll 2007; Edmunds, Leichter, and Adjeroud 2010), it is possible that: 1) *P. meandrina* also spawns around the full moons of January and February, 2) that pelagic larval duration (PLD) can extend over 1-2 months; and/or 3) that connected populations of *Pocillopora meandrina* (e.g., throughout French Polynesia) may spawn at different times, or PLD varies in those populations. Alternatively, the recruitment data of *Pocillopora* spp. could represent multiple species spawning across different months, and have only been identified to genus on the recruitment tiles. These competing hypotheses identify high priorities for further research as marine heatwaves both drive mass mortality (Hughes, Kerry, and Simpson 2018; Speare et al. 2022) and also can impact reproductive timing (T. Shlesinger and Loya 2019), amount/quality (Johnston et al. 2020; A. H. Baird and Marshall 2002), and recruitment (Peter J. Edmunds 2017; Hughes et al. 2019).

### Pocillopora grandis

Given the complexity of identification, very little observational literature on spawning in *P. grandis* (Previously named *P. eydouxi*) has been recorded *(Smith et al. 2008; Schmidt-Roach et al. 2012; Hirose, Kinzie, and Hidaka 2000)*. The first observations with visual identification in Okinawa Japan made by (Hirose, Kinzie, and Hidaka 2000), indicated spawning a few days after the full moon, which is in line with our findings. A second study on the Great Barrier Reef in Australia identified *P. grandis* both visually and by mtORF haplotype (Schmidt-Roach et al. 2012), but not to the species level with PocHistone (Johnston, Forsman, and Toonen 2018). The Australia study reported spawning on full moon and full moon +1, which matches our observations in Mo’orea (Fig. 4). However, in Australia both *P. meandrina* and *P. grandis* were reported with the same diel timing (06:25), whereas we documented distinctly different diel timing, with *P. grandis* spawning first and *P. meandrina* 25 minutes later (Fig. 5). This paucity of data across the broad geographic distribution of *P. grandis* indicates further study is necessary in multiple areas of the world to enable capacity to complete location specific differences, and their proximate environmental causes, in lunar and diel timing.

### Pocillopora verrucosa

*P. verrucosa* in Mo’orea only spawned in association with the new moon on lunar day 0 and 1 . Similarly, in the Red Sea (Sudan region), morphologically identified *P. verrucosa* spawning was observed to take place -1 day before the new moon, and in Saudi Arabia at -2 days before the new moon (Bouwmeester et al. 2021; Bouwmeester, Berumen, and Baird 2011; Bouwmeester et al. 2015). Conversely, morphologically identified *P. verrucosa* spawning has been reported on full moon lunar day 0 and 1 on the GBR (Schmidt-Roach et al. 2012) and 1–3 days following the full moon in Lyudao Taiwan (Lin and Nozawa 2017; Mulla et al. 2021). These reports are outliers from the broader geographical pattern we are seeing in Mo’orea and the reports of new moon morphologically identified *P. verrucosa* spawning in the Red Sea (Bouwmeester, Berumen, and Baird 2011; Bouwmeester et al. 2021; Bouwmeester et al. 2015), suggest species differences are present.

When considering diel timing of release, in Mo’orea (17.53S latitude), *P. verrucosa* gametes were released ∼5:45-6:20 local time, ∼38 to 55 min after sunrise local time, closer to sunrise than morphologically identified *P. verrucosa* in several other regions of the world (Table 1). These findings suggest two hypotheses. First, that there are latitudinal differences in diel/lunar spawning times and second that there are different cryptic *Pocillopora spp.* that can’t be compared in the absence of genetic identification in other studies. While spawning of morphologically identified *P. verrucosa* at the island of Okinawa, Japan (26.60N latitude) occurs within the same time range as Mo’orea at ∼1h after sunrise (Kinzie 1993, Hirose et al. 2000), morphologically identified *P. verrucosa* closer to the equator at the island of Ishigaki, Japan (24.42N latitude) was observed to spawn ∼3h after sunrise (Suzuki 2012). Similarly to Ishigaki Japan, in the Red Sea morphologically identified *P. verrucosa* spawning has been documented at ∼3h after sunrise (∼08:45) on reefs in Bara Musa Saqir Reef in Sudan (19.05N latitude) and Al Jadir Reef in Saudi Arabia (19.79 latitude) (Bouwmeester et al. 2021). The same diel timing of morphologically identified *P. verrucosa* spawning (∼08:40 - 09:20) was also documented in the central Red Sea Thuwal, (22.22N latitude)(JBouwmeester, Berumen, and Baird 2011) and Al Fahal (22.22N latitude) and Dreams Beach Reef (21.76 latitude) (Bouwmeester et al. 2015). Lastly, morphologically identified *P. verrucosa* in Lyudao Taiwan (22.66N latitude), the reports of ∼5 h after sunrise (09:30 to 10:30) (Lin and Nozawa 2017; Mulla et al. 2021). Peak spawning time occurring later in the season further from the equator has been documented in multiple corals (Howells et al. 2014), but the lunar data mismatch of our genetically identified *P. verrucosa* as new moon and two other morphologically identified *P. verrucosa* as full moon spawners (Schmidt-Roach et al. 2012; Lin and Nozawa 2017), suggests species identification could explain these different patterns instead.

Previously morphologically identified *P. verrucosa* in the Red Sea is actually *P. favosa* (Oury et al. 2025). The identification of *P. favosa* could explain the several hour difference in time of gamete release (<2h after sunrise in Mo’oera, versus 3 - 4 hours after sunrise in the Red Sea) and indicates that no global spawning conclusions should be made for *P. verrucosa* due to the genetic differences within the morphological group. Further, not every coral observed spawning in the GBR report (Schmidt-Roach et al. 2012) was identified genetically, and none of the corals in the Taiwan observations (Lin and Nozawa 2017; Mulla et al. 2021) were identified genetically. Therefore it is highly likely that contrasting results by reef location are due to species identity. Therefore we propose a broad geographic distribution of *P. verrucosa* spawning associated with the new moon based on genetic species identification. Despite the greater amount of studies of spawning in *P. verrucosa*, comparisons of latitudinal drivers can not yet be made in the absence of genetic identification. In the future latitudinal associated drivers in spawn timing (e.g., (Howells et al. 2014)) should also be tested in spawning *Pocillopora spp.* It is essential that studies identify *Pocillopora* based on now-established genetic markers (Johnston and Burgess 2023) and/or genomic data (Johnston, Cunning, and Burgess 2022).

### Pocillopora tuahiniensis

At the same site where we observed the spawning of *P. meandrina* and *P. verrucosa* species, we also observed the spawning of *Pocillopora tuahiniensis (Johnston and Burgess 2023)*. *P. verrucosa* and *P. tuahiniensis* are sister species and overlap in their distribution at Mo’orea, across habitats and depth (Johnston et al. 2022). *P. verrucosa* has a much broader geographic distribution across the Indo-Pacific (Gelin 2017 and Oury 2023 refs) compared to *P. tuahiniensis*; the latter has been documented within a subset of the range of the former (French Polynesia (Oury, Gélin, and Magalon 2021; Edmunds et al. 2016; Gélin et al. 2017; Forsman et al. 2013)), Ducie Island, and in Rapa Nui (Armstrong et al. 2023; Voolstra et al. 2023). The close phylogenetic relationship between *P. tuahiniensis* and *P. verrucosa* and their overlap in geographic ranges and distributions within reefs, strongly suggests that the temporal separation in gamete release, in terms of both the day after new moon and the minutes after sunrise on overlapping lunar days, provides a mechanism to maintain species boundaries (preventing gene flow) between these species.

In our observations, *P. verrucosa* spawning (∼05:55 - 06:20) was temporally offset from its sister species *P. tuahiniensis* (∼06:30 - 06:50) by ∼28 minutes. Our results are consistent with a linkage between genetic divergence and the temporal divergence in diel spawning time we observed here. Further, *Pocillopora* spp. embryo development is some of the most rapid of corals described to date, where fertilization can occur within 15 minutes (Mulla et al. 2021), cleavage within 30 - 40 minutes, and full development to the swimming larval stage by 8 hours post fertilization (Hirose, Kinzie, and Hidaka 2000). Given this rapid fertilization and cleavage and the quick dissipation of the clouds of egg and sperm following spawning, the ∼30 minute offset between spawning in these two species appears to be one line of evidence that would minimize the potential for cross fertilization between species. Genome-wide analyses of these species show no evidence of hybridization (Johnston, Cunning, and Burgess 2022), further supporting that the timing offset of spawning between *P. verrucosa* and *P. tuahiniensis* is acting as a temporal barrier for speciation.

### Pocillopora effusa

*P. effusa* is less abundant in the lagoon than on the fore reef (Burgess, Turner, and Johnston 2024). In addition, this species recently suffered a significant loss of abundance due to bleaching at Mo’orea in 2019 (Burgess et al. 2021). Within this context, observations of *P. effusa* in the lagoon are likely to be rare. Indeed, only a single colony of *P. effusa* was observed spawning on lunar day 3 after the November 2023 new moon (15 November) at ∼06:31-06:50. Because this was a single observation, it could indicate that spawning is occurring at different times when observers were not present, or multiple months as we observed with the other *Pocillopora* species in our study. Alternatively, it could be due to low abundance of colonies at our observation site, or accidental assignment of adjacent colony spawning to *P. effusa*. Therefore, it is necessary to conduct additional sampling with extended time of day, days in a month, and additional months in future studies to concretely assign the timing of spawning for *P. effusa*.

### Larvae contain zooxanthellae and development is rapid

For all *Pocillopora* species we observed, once spawning commenced, the water quickly became cloudy with eggs and sperm (e.g., Fig. 1A, B, C). Eggs appeared to be neutrally or negatively buoyant within collection bottles, which has been reported by (Suzuki 2012; Schmidt-Roach et al. 2012; Mulla et al. 2021), as we observed they did not rise quickly to the surface like the larger, lipid rich egg bundles of Acroporidae. Following release, the gametes were rapidly dispersed by the waves and current. From our collections, eggs (∼90µm) and developing embryos (∼120µm) that were observed at ∼10:00 on the day of release via compound microscopy contained zooxanthellae, which have been shown to enter the eggs in the ∼6 days prior to spawning in *Pocillopora* in Japan (Hirose, Kinzie, and Hidaka 2001). By 14:00 (∼7.5 hours post fertilization, hpf) larvae were actively swimming, matching the timing of development in the morphologically identified *P. verrucosa* and *P. eydouxi* in Japan, where ciliated swimming larvae were documented 8 hpf (Hirose, Kinzie, and Hidaka 2000). The rapid development of Pocilloporidae larvae with vertically transmitted Symbiodiniaceae that can provide nutritional supplementation (Kopp et al. 2016; Huffmyer et al. 2025) could affect recruitment success (Adjeroud, Penin, and Carroll 2007; Gleason 1996; Edmunds, Leichter, and Adjeroud 2010). Rapid development of symbiotic larvae could influence the capacity of *Pocillopora* spp. to promote rapid recovery following disturbances on the reefs of Mo’orea, provided there is enough adult colonies (Speare et al. 2022) and that gametogenesis is not severely impaired by heatwaves and mass bleaching events (Johnston et al. 2020).

Further work on the clarification of the diel timing of spawning, development, and settlement of spawned Pocilloporid symbiotic larvae will be essential. Both spawning (Keith et al. 2016; Baird, Guest, and Willis 2009) and development (Woolsey, Byrne, and Baird 2013; Negri, Marshall, and Heyward 2007; Randall and Szmant 2009) are temperature dependent. Additionally, symbiotic larvae are light dependent with respect to metabolism (Kopp et al. 2016; Huffmyer et al. 2023) and phototaxis swimming (Mulla et al. 2021), resulting in the potential for UV damage to host and symbiont. For example, embryo developmental abnormalities increase with increasing temperature (Bassim, Sammarco, and Snell 2002) and symbiotic larvae accumulate more DNA damage from UV than aposymbiotic larvae (Baird et al. 2021; Nesa et al. 2012). Therefore understanding the diel timing of release will be essential for quantifying the amount of irradiance and diel solar warming *Pocillopora* larvae will experience in the critical first 8 hours of development to the swimming stage.

*Pocillopora spp.* can dominate coral reefs (Pérez-Rosales et al. 2021) and facilitate rapid reef recovery (Holbrook et al. 2018). Cryptic *Pocillopora* species, however, exhibit differences in their thermal sensitivity (Burgess et al. 2021). Here, we have documented differences in their timing of spawning in terms of month, lunar phase, and time of day. Future work can now build on these findings to determine key aspects such as thermal tolerance of larval development and early life stages, pelagic larval duration, and settlement preferences, which will improve ecological forecasting capabilities for tropical reefs throughout the Indo-Pacific.

## Acknowledgements

As guests, we recognize and give thanks for the land and water resources of French Polynesia, in particular Mo’orea, and to the traditional owners of the land, both past and present for the resources studied in this work. Māuruuru roa. With respect to the spelling of Tahitian words, we endeavored to follow the Te Fare Vāna’a transcription system that is adhered to by a large segment of the Tahitian community, but also recognize other community members follow the Raapoto transcription system where the island name of Moorea is, for example, spelled without the ’eta (i.e., Moorea). The samples collected herein are resources of the people and government of French Polynesia and have been supplied under the Direction de l’environnement DIREN Permit Number 10626 (06 November 2023) and exported under Convention on International Trade in Endangered Species CITES export permit FR24987000029-E.

Although we are the first to report scientific documentation of *Pocillopora meandrina, P. verrucosa*, *P. tuahiniensis* and *P. grandis* spawning in French Polynesia, we also want to acknowledge the likelihood that that the timing of coral spawning for multiple taxa has been known by local Polynesians for decades to centuries. Merging of traditional ecological knowledge, citizen science, and scientific monitoring and documentation is poised to greatly expand our understanding of reproduction in French Polynesia, which will be critical to reef recovery and maintenance under climate change.

En tant qu’invités, nous reconnaissons et exprimons notre gratitude envers les ressources terrestres et aquatiques de la Polynésie française, en particulier de Mo’orea, ainsi qu’envers les propriétaires traditionnels de ces terres, passés et presents pour les ressources étudiées dans ce travail. Māuruuru roa. En ce qui concerne l’orthographe des mots tahitiens, nous avons fait notre possible pour suivre le système de transcription Te Fare Vāna’a, auquel adhère une grande partie de la communauté tahitienne, tout en reconnaissant que d’autres membres de la communauté suivent le système de transcription Raapoto où le nom de l’île de Moorea, par exemple, est orthographié sans le ’eta (c’est-à-dire Moorea). Les échantillons collectés ici sont des ressources du peuple et du gouvernement de la Polynésie française et ont été fournis sous le permis de la Direction de l’environnement DIREN numéro 10626 (06 novembre 2023) et exportés sous le permis d’exportation de la Convention sur le commerce international des espèces de faune et de flore sauvages menacées d’extinction CITES FR24987000029-E.

Bien que nous soyons les premiers à rapporter une documentation scientifique sur la ponte de *Pocillopora meandrina*, *P. verrucosa*, *P. tuahiniensis* et *P. grandis* en Polynésie française, nous tenons également à reconnaître qu’il est probable que le calendrier de reproduction de plusieurs taxons de coraux soit connu des Polynésiens depuis des décennies, voire des siècles. La combinaison des savoirs écologiques traditionnels, de la science participative et du suivi et de la documentation scientifiques est en passe d’élargir considérablement notre compréhension de la reproduction en Polynésie française, ce qui sera essentiel pour la restauration et le maintien des récifs face au changement climatique.

We are thankful for observer support from Ariana Huffmyer, Danielle Becker, Chloé Gilligan, Erica Atkins, Danielle Barnes, Holly Moeller, Arnaud Fabregues, Guillaume Iwankow, Lucy Gorman, Ross Cunning, Shayle Matsuda, Tali Mass, Keisha Bahr, Cody Clements, Kelly Speare, and Raine Detmer and for the staff of the UC Richard B. Gump South Pacific Research Station for facilitating our research. We appreciate discussions with and feedback from Peter Edmunds and Ariana Huffmyer. This research was funded by the U.S. National Science Foundation Grants EF-1921465 and OCE-2348674 to HMP and OCE-1829867 to SCB and supported by resources from NSF-OCE 2224354 to the Mo’orea Coral Reef LTER, as well as a generous gift from the Gordon and Betty Moore Foundation.

## Data Availability

DOI 10.17605/OSF.IO/5GEQD (Harnay et al. 2023)

## Author Contributions

Funding HMP, SCB; Conceptualization PEH, HMP; Field work PEH, HMP; Molecular Analysis PEH, HMP, AT, SCB; Writing PEH, HMP, AT, SCB

## Supplementary material

**Table S1.** *Pocillopora* spp. spawning observation timeline from September 2022 to December 2024. Moon phase dates were reported based on the website https://aa.usno.navy.mil/data/MoonPhases. Sunrise and sunset times were reported for the location of Mo’orea using this website https://www.timeanddate.com/sun/@4034185 NDR: <u>N</u>o <u>D</u>ata <u>R</u>ecorded indicates no observers were present in the field.

- - - indicates no additional notes

** Indicates species were identified from previously tagged and sequenced colonies

**Table S2.** *Pocillopora* spawning sample numbers and genetic species identification collected from corals observed spawning in 2022, 2023, and 2024.

