## Supplementary material for "Daytime broadcast spawning in cryptic *Pocillopora* species at Mo’orea, French Polynesia"

Data, Code, Image Archive: <https://osf.io/5geqd/>

**Abstract**

Knowledge of coral gamete release is fundamental to reef ecology and evolution, but cryptic species complicate reproduction studies. In Mo’orea, French Polynesia, when corals were decimated by crown-of-thorns and a cyclone between 2007-2010, recruitment of six cryptic species of *Pocillopora spp.* primarily drove decadal reef recovery. Broadcast spawning for these genetically-verified species at Mo’orea is, however, undocumented in the scientific literature. We conducted *in situ* surveys over 58 days during September 2022-January 2023, October 2023-January 2024, and October 2024-December 2024. To identify the *Pocillopora spp.* we used molecular analysis of mtORF and PocHistone markers. *P. meandrina* and *P. grandis* spawned 2-3 days following the full moons of October-December. In contrast, *P. verrucosa* and *P. tuahiniensis* spawned at the new moons in November-December, with *P. verrucosa* spawning on lunar days 0-1, and *P. tuahiniensis* spawning on lunar days 1-3. Diel offset in the timing of spawning following sunrise was present in both moon phases indicating a temporal speciation barrier. *P. verrucosa* spawned ~45 min after sunrise, whereas *P. tuahiniensis* spawned ~73 min after sunrise. Similarly, *P. grandis* spawned ~40 minutes after sunrise, whereas *P. meandrina* spawned ~65 minutes after sunrise. Only one colony of *P. effusa* spawned (lunar day 3 new moon ~06:30-06:50), requiring further confirmation of the timing for this low abundance species. Collectively, these first spawning reports provide an initial scientific documentation of *Pocillopora* spawning at the genetically-verified species level in Mo’orea, increasing capacity to study the essential process of coral reproduction for critical reef building species.

**Supplementary material:**

**Table S1.** *Pocillopora* spp. spawning observation timeline from September 2022 to December 2024. Moon phase dates were reported based on the website <https://aa.usno.navy.mil/data/MoonPhases>. Sunrise and sunset times were reported for the location of Mo’orea using this website <https://www.timeanddate.com/sun/@4034185>

NDR: No Data Recorded indicates no observers were present in the field.

- - - indicates no additional notes

** Indicates species were identified from previously tagged and sequenced colonies

| **Date**  **YYYYMMDD** | **Sunrise Time** | **Lunar Phase** | **Spawn** | **Observation Start Time** | **Observation Stop Time** | **Spawning Timing and Notes** |
| --- | --- | --- | --- | --- | --- | --- |
| 20220925 | 5:46 AM | New | NDR | NDR | NDR | - - - |
| 20220926 | 5:45 AM | New+1 | No | 05:30 AM | 7:00 AM | - - - |
| 20220927 | 5:44 AM | New+2 | No | 05:40 AM | 8:00 AM | - - - |
| 20220928 | 5:43 AM | New+3 | No | 05:45 AM | 08:15 AM | - - - |
| 20221007 | 5:36 AM | Full-2 | No | 05:45 AM | 08:00 AM | - - - |
| 20221008 | 5:35 AM | Full-1 | No | 06:00 AM | 08:05 AM | - - - |
| 20221009 | 5:34 AM | Full | NDR | NDR | NDR | - - - |
| 20221010 | 5:34 AM | Full +1 | NDR | NDR | NDR | - - - |
| 20221011 | 5:33 AM | Full +2 | NDR | NDR | NDR | - - - |
| 20221012 | 5:32 AM | Full +3 | NDR | NDR | NDR | - - - |
| 20221106 | 5:18 AM | Full-2 | No | 6:00 AM | 7:45 AM | - - - |
| 20221107 | 5:17 AM | Full-1 | No | 6:10 AM | 8:00 AM | - - - |
| 20221108 | 5:17 AM | Full | NDR | NDR | NDR | Bad weather with lightning precluded in water observations |
| 20221109 | 5:15 AM | Full+1 | No | 5:50 AM | 8:00 AM | - - - |
| 20221110 | 5:15 AM | Full+2 | Yes | 05:45 AM | 08:25 AM | 06:32 - unknown  *P. meandrina* |
| 20221111 | 5:14 AM | Full+3 | No | 6:00 AM | 7:40 AM | - - - |
| 20221205 | 5:16 AM | Full -2 | NDR | NDR | NDR | - - - |
| 20221206 | 5:16 AM | Full -1 | NDR | NDR | NDR | - - - |
| 20221207 | 5:16 AM | Full | No | 06:00 AM | 07:28 AM | - - - |
| 20221208 | 5:16 AM | Full+1 | No | 06:20 AM | 07:52 AM | - - - |
| 20221209 | 5:17 AM | Full+2 | Yes | 06:05 AM | 08:00 AM | 06:16 - 06:32  *P. meandrina* |
| 20221210 | 5:17 AM | Full+3 | Yes | 06:00 AM | 07:45 AM | 06:21 - 06:35  *P. meandrina* |
| 20230104 | 5:29 AM | Full -2 | NDR | NDR | NDR | - - - |
| 20230105 | 5:30 AM | Full -1 | NDR | NDR | NDR | - - - |
| 20230106 | 5:29 AM | Full | NDR | NDR | NDR | Bad weather with lightning precluded in water observations |
| 20230107 | 5:30 AM | Full+1 | No | 06:09 AM | 07:37 AM | Bad weather and strong current reduced in water time |
| 20230108 | 5:30 AM | Full+2 | No | 06:10 AM | 07:18 AM | Bad weather and strong current reduced in water time |
| 20230109 | 5:33 AM | Full+3 | No | 06:10 AM | 06:37 AM | Bad weather, waves and current precluded water work past ~06:40 |
| 20230110 | 5:33 AM | Full +4 | NDR | NDR | NDR | - - - |
| 20230111 | 5:34 AM | Full +5 | NDR | NDR | NDR | - - - |
| 20231026 | 5:23 AM | Full -2 | NDR | NDR | NDR | - - - |
| 20231027 | 5:23 AM | Full -1 | NDR | NDR | NDR | - - - |
| 20231028 | 5:22 AM | Full | No | 6:00 AM | 7:30 AM | - - - |
| 20231029 | 5:22 AM | Full +1 | No | 6:07 AM | 8:00 AM | - - - |
| 20231030 | 5:21 AM | Full +2 | Yes | 6:00 AM | 8:00 AM | 06:22 - 06:38  *P. meandrina* |
| 20231031 | 5:21 AM | Full +3 | No | 6:08 AM | 8:02 AM | - - - |
| 20231101 | 5:20 AM | Full +4 | No | 6:18 AM | 7:40 AM | - - - |
| 20231102 | 5:20 AM | Full +5 | No | 6:04 AM | 7:12 AM | - - - |
| 20231103 | 5:19 AM | Full +6 | No | 6:10 AM | 7:30 AM | Low visibility |
| 20231104 | 5:19 AM | Full +7 | No | 6:02 AM | 7:30 AM | Low visibility |
| 20231110 | 5:17 AM | New -2 | NDR | NDR | NDR | - - - |
| 20231111 | 5:17 AM | New -1 | NDR | NDR | NDR | - - - |
| 20231112 | 5:16 AM | New | Yes | 5:52 AM | 7:30 AM | 06:00 - 06:17  *P. verrucosa* |
| 20231113 | 5:16 AM | New +1 | Yes | 5:49 AM | 7:08 AM | 6:04 - 6:19  *P. verrucosa* |
| 20231113 | 5:16 AM | New +1 | Yes | 5:49 AM | 7:08 AM | 6:33 - 6:44  *P. tuahiniensis* |
| 20231114 | 5:16 AM | New +2 | Yes | 5:50 AM | 7:18 AM | 6:26 - 6:50  *P. tuahiniensis* |
| 20231115 | 5:15 AM | New +3 | Yes | 5:55 AM | 7:30 AM | 6:31 - 6:50  *P. tuahiniensis*  *P. effusa* |
| 20231116 | 5:15 AM | New +4 | No | 5:54 AM | 7:24 AM | lot of wind and wave at the water surface but good visibility |
| 20231117 | 5:15 AM | New +5 | No | 5:57 AM | 7:04 AM | lot of wind |
| 20231124 | 5:14 AM | Full -2 | No | 6:00 AM | 7:00 AM | - - - |
| 20231125 | 5:14 AM | Full -1 | No | 5:50 AM | 7:10 AM | - - - |
| 20231126 | 5:14 AM | Full | No | 5:49 AM | 7:09 AM | - - - |
| 20231127 | 5:15 AM | Full +1 | NDR | NDR | NDR | Big swell, high current. Visibility zero |
| 20231128 | 5:15 AM | Full +2 | NDR | NDR | NDR | - - - |
| 20231129 | 5:15 AM | Full +3 | NDR | NDR | NDR | - - - |
| 20231130 | 5:15 AM | Full +4 | NDR | NDR | NDR | - - - |
| 20231201 | 5:15 AM | Full +5 | NDR | NDR | NDR | - - - |
| 20231210 | 5:17 AM | New -2 | No | 6:02 AM | 6:57 AM | - - - |
| 20231211 | 5:17 AM | New -1 | No | 6:06 AM | 7:03 AM | - - - |
| 20231212 | 5:18 AM | New | Yes | 5:45 AM | 7:00 AM | 5:56 - 6:17  *P. verrucosa* |
| 20231213 | 5:18 AM | New +1 | NDR | NDR | NDR | Huge Storm, too dangerous to monitor |
| 20231214 | 5:18 AM | New +2 | Yes | 6:06 AM | 7:05 AM | 6:33 - 6:42  *P. tuahiniensis* |
| 20231215 | 5:19 AM | New +3 | Yes | 6:06 AM | 7:00 AM | 6:34 - 6:36  No colony genetically identified |
| 20231216 | 5:19 AM | New +4 | NDR | NDR | NDR | - - - |
| 20231217 | 5:20 AM | New +5 | NDR | NDR | NDR | - - - |
| 20231224 | 5:23 AM | Full -2 | NDR | NDR | NDR | - - - |
| 20231225 | 5:24 AM | Full -1 | NDR | NDR | NDR | - - - |
| 20231226 | 5:24 AM | Full | No | 6:00 AM | 7:05 AM | - - - |
| 20231227 | 5:25 AM | Full +1 | No | 6:01 AM | 7:35 AM | - - - |
| 20231228 | 5:25 AM | Full +2 | NDR | NDR | NDR | - - - |
| 20231229 | 5:26 AM | Full +3 | NDR | NDR | NDR | - - - |
| 20231230 | 5:26 AM | Full +4 | NDR | NDR | NDR | - - - |
| 20231231 | 5:27 AM | Full +5 | NDR | NDR | NDR | - - - |
| 20240109 | 5:33 AM | New -2 | NDR | NDR | NDR | - - - |
| 20240110 | 5:33 AM | New -1 | NDR | NDR | NDR | - - - |
| 20240111 | 5:34 AM | New | NDR | NDR | NDR | - - - |
| 20240112 | 5:34 AM | Newl +1 | No | 6:05 AM | 7:05 AM | Strong current, the day before big storm and rain |
| 20240113 | 5:35 AM | New +2 | No | 5:45 AM | 7:05 AM | Strong current |
| 20240114 | 5:36 AM | New +3 | No | 5:40 AM | 7:00 AM | Strong current |
| 20240115 | 5:36 AM | New +4 | NDR | NDR | NDR | - - - |
| 20240116 | 5:37 AM | New +5 | NDR | NDR | NDR | - - - |
| 20240123 | 5:41 AM | Full -2 | NDR | NDR | NDR | - - - |
| 20240124 | 5:42 AM | Full -1 | NDR | NDR | NDR | - - - |
| 20240125 | 5:42 AM | Full | No | 6:02 AM | 6:15 AM | High wind impossible to monitor longer because of the swell and the current |
| 20240126 | 5:43 AM | Full +1 | No | 5:55 AM | 7:05 AM | - - - |
| 20240127 | 5:44 AM | Full +2 | No | 5:50 AM | 7:10 AM | - - - |
| 20240128 | 5:44 AM | Full +3 | No | 5:50 AM | 7:10 AM | - - - |
| 20240129 | 5:45 AM | Full +4 | NDR | NDR | NDR | - - - |
| 20240130 | 5:45 AM | Full +5 | NDR | NDR | NDR | - - - |
| 20241101 | 5:20 AM | New | Yes | 5:45 AM | 7:05 AM | 6:15 - 6:25  *P. verrucosa* ** |
| 20241102 | 5:19 AM | New+1 | Yes | 5:49 AM | 7:05 AM | 6:35 - 6:58  *P. tuahiniensis* ** |
| 20241103 | 5:19 AM | New+2 | Yes | 5:55 AM | 7:05 AM | 6:30 - 6:55  *P. tuahiniensis* ** |
| 20241117 | 5:14 AM | Full +2 | Yes | 5:45 AM | 7:00 AM | 5:55 - 6:11  *P. grandis* |
| 20241117 | 5:14 AM | Full +2 | Yes | 5:45 AM | 7:00 AM | 6:23 - 6:37  *P. meandrina* ** |
| 20241118 | 5:14 AM | Full +3 | Yes | 5:45 AM | 7:05 AM | 5:56 - 6:05  *P. grandis* |
| 20241118 | 5:14 AM | Full +3 | Yes | 5:45 AM | 7:05 AM | 6:18 - 6:28  *P. meandrina* ** |
| 20241130 | 5:15 AM | New | Yes | 5:45 AM | 7:00 AM | 05:56 - 06:15  *P. verrucosa* ** |
| 20241201 | 5:15 AM | New +1 | Yes | 5:45 AM | 7:00 AM | 05:58-unknown  *P. verrucosa* ** |
| 20241201 | 5:15 AM | New +1 | Yes | 5:45 AM | 7:00 AM | 06:26-unknown  *P. tuahiniensis* ** |
| 20241202 | 5:15 AM | New +2 | Yes | 5:55 AM | 7:05 AM | 06:19 - 06:46  *P. tuahiniensis* ** |
| 20241216 | 5:19 AM | Full +2 | Yes | 5:35AM | 7:15 AM | 6:01 - 6:12  *P. grandis* |
| 20241216 | 5:19 AM | Full +2 | Yes | 5:35AM | 7:15 AM | 6:23 - 6:31  *P. meandrina* ** |
| 20241217 | 5:19 AM | Full +3 | Yes | 5:40 AM | 7:05 AM | 5:55 - 6:10  *P. grandis* |
| 20241217 | 5:19 AM | Full +3 | Yes | 5:40 AM | 7:05 AM | 06:21 - 06:32  *P. meandrina* ** |

**Table S2.** *Pocillopora* spawning samples collected from corals observed spawning in 2022, 2023, and 2024.

| **Sample ID** | **mtORF Sanger ID**  **Haplotype** | **PocHistone RFLP ID** | **Species ID** | **Collection Date**  **YYYYMMDD** |
| --- | --- | --- | --- | --- |
| POC-578 (Tag 192) | Haplotype 1 | *P. meandrina* | *P. meandrina* | 20221110 |
| POC-131 | Haplotype 1 | *P. meandrina* | *P. meandrina* | 20221209 |
| POC-132 | Haplotype 1 | *P. meandrina* | *P. meandrina* | 20221209 |
| POC-133 | Haplotype 8 | *NA* | *P. meandrina* | 20221209 |
| POC-134 | Haplotype 8 | *NA* | *P. meandrina* | 20221209 |
| POC-135 | Haplotype 1 | *P. meandrina* | *P. meandrina* | 20221209 |
| POC-176 | Haplotype 1 | *P. meandrina* | *P. meandrina* | 20221209 |
| POC-185 | Haplotype 1 | *P. meandrina* | *P. meandrina* | 20221209 |
| POC-186 | Haplotype 1 | *P. meandrina* | *P. meandrina* | 20221209 |
| POC-189 | Haplotype 1 | *P. meandrina* | *P. meandrina* | 20221209 |
| POC-191 | Haplotype 1 | *P. meandrina* | *P. meandrina* | 20221209 |
| POC-194 | Haplotype 1 | *P. meandrina* | *P. meandrina* | 20221209 |
| POC-195 | Haplotype 1 | *P. meandrina* | *P. meandrina* | 20221209 |
| POC-196 | Haplotype 1 | *P. meandrina* | *P. meandrina* | 20221209 |
| POC-199 | Haplotype 1 | *P. meandrina* | *P. meandrina* | 20221209 |
| POC-200 | Haplotype 8 | *NA* | *P. meandrina* | 20221209 |
| POC-002 | Haplotype 1 | *P. meandrina* | *P. meandrina* | 20231030 |
| POC-001 | Haplotype 1 | *P. meandrina* | *P. meandrina* | 20231030 |
| POC-003 | Haplotype 1 | *P. meandrina* | *P. meandrina* | 20231030 |
| POC-011 | Haplotype 3 | *NA* | *P. verrucosa* | 20231112 |
| POC-012 | Haplotype 3 | *NA* | *P. verrucosa* | 20231112 |
| POC-014 | Haplotype 3 | *NA* | *P. verrucosa* | 20231112 |
| POC-017 | Haplotype 3 | *NA* | *P. verrucosa* | 20231112 |
| POC-019 | Haplotype 3 | *NA* | *P. verrucosa* | 20231112 |
| POC-020 | Haplotype 3 | *NA* | *P. verrucosa* | 20231112 |
| POC-004 | Haplotype 3 | *NA* | *P. verrucosa* | 20231112 |
| POC-005 | Haplotype 3 | *NA* | *P. verrucosa* | 20231112 |
| POC-022 | Haplotype 10 | *NA* | *P. tuahiniensis* | 20231113 |
| POC-047 | Haplotype 10 | *NA* | *P. tuahiniensis* | 20231113 |
| POC-007 | Haplotype 3 | *NA* | *P. verrucosa* | 20231113 |
| POC-021 | Haplotype 3 | *NA* | *P. verrucosa* | 20231113 |
| POC-028 | Haplotype 3 | *NA* | *P. verrucosa* | 20231113 |
| POC-041 | Haplotype 3 | *NA* | *P. verrucosa* | 20231113 |
| POC-042 | Haplotype 3 | *NA* | *P. verrucosa* | 20231113 |
| POC-006 | Haplotype 3 | *NA* | *P. verrucosa* | 20231113 |
| POC-015 | Haplotype 10 | *NA* | *P. tuahiniensis* | 20231114 |
| POC-023 | Haplotype 10 | *NA* | *P. tuahiniensis* | 20231114 |
| POC-024 | Haplotype 10 | *NA* | *P. tuahiniensis* | 20231114 |
| POC-025 | Haplotype 10 | *NA* | *P. tuahiniensis* | 20231114 |
| POC-026 | Haplotype 10 | *NA* | *P. tuahiniensis* | 20231114 |
| POC-029 | Haplotype 10 | *NA* | *P. tuahiniensis* | 20231114 |
| POC-030 | Haplotype 10 | *NA* | *P. tuahiniensis* | 20231114 |
| POC-033 | Haplotype 10 | *NA* | *P. tuahiniensis* | 20231114 |
| POC-038 | Haplotype 10 | *NA* | *P. tuahiniensis* | 20231114 |
| POC-045 | Haplotype 10 | *NA* | *P. tuahiniensis* | 20231114 |
| POC-046 | Haplotype 10 | *NA* | *P. tuahiniensis* | 20231114 |
| POC-048 | Haplotype 10 | *NA* | *P. tuahiniensis* | 20231114 |
| POC-050 | Haplotype 10 | *NA* | *P. tuahiniensis* | 20231114 |
| POC-051 | Haplotype 10 | *NA* | *P. tuahiniensis* | 20231114 |
| POC-009 | Haplotype 10 | *NA* | *P. tuahiniensis* | 20231114 |
| POC-010 | Haplotype 10 | *NA* | *P. tuahiniensis* | 20231114 |
| POC-180 | Haplotype 10 | *NA* | *P. tuahiniensis* | 20231114 |
| POC-044 | Haplotype 11 | *NA* | *P.* *effusa* | 20231115 |
| POC-034 | Haplotype 10 | *NA* | *P. tuahiniensis* | 20231115 |
| POC-035 | Haplotype 10 | *NA* | *P. tuahiniensis* | 20231115 |
| POC-036 | Haplotype 10 | *NA* | *P. tuahiniensis* | 20231115 |
| POC-037 | Haplotype 10 | *NA* | *P. tuahiniensis* | 20231115 |
| POC-039 | Haplotype 10 | *NA* | *P. tuahiniensis* | 20231115 |
| POC-040 | Haplotype 10 | *NA* | *P. tuahiniensis* | 20231115 |
| POC-043 | Haplotype 10 | *NA* | *P. tuahiniensis* | 20231115 |
| POC-049 | Haplotype 10 | *NA* | *P. tuahiniensis* | 20231115 |
| POC-052 | Haplotype 10 | *NA* | *P. tuahiniensis* | 20231115 |
| POC-053 | Haplotype 3 | *NA* | *P. verrucosa* | 20231212 |
| POC-055 | Haplotype 3 | *NA* | *P. verrucosa* | 20231212 |
| POC-057 | Haplotype 3 | *NA* | *P. verrucosa* | 20231212 |
| POC-060 | Haplotype 3 | *NA* | *P. verrucosa* | 20231212 |
| POC-061 | Haplotype 3 | *NA* | *P. verrucosa* | 20231212 |
| POC-069 | Haplotype 3 | *NA* | *P. verrucosa* | 20231212 |
| POC-070 | Haplotype 3 | *NA* | *P. verrucosa* | 20231212 |
| POC-182 | Haplotype 3 | *NA* | *P. verrucosa* | 20231212 |
| POC-090 | Haplotype 10 | *NA* | *P. tuahiniensis* | 20231214 |
| POC-074 | Haplotype 10 | *NA* | *P. tuahiniensis* | 20231214 |
| POC-151 | Haplotype 1 | *P. grandis* | *P. grandis* | 20241117 |
| POC-178 | Haplotype 1 | *P. grandis* | *P. grandis* | 20241117 |
| POC-150 | Haplotype 1 | *P. grandis* | *P. grandis* | 20241117 |
